# *In vitro* activity of combination formulations of the novel metallo-β-lactamase (MBL) inhibitor APC148 with comparator treatments against 176 MBL-containing Enterobacterales isolates from the SENTRY Antimicrobial Surveillance Program (2019-2022)

**DOI:** 10.64898/2026.03.12.711254

**Authors:** Veronika Smith, Bjørn Klem, Bjørg Bolstad, Hanne Cecilie Winther-Larsen, Ole Andreas Løchen Økstad, Pål Rongved

**Author notes:** Address correspondence to Pål Rongved.

## Abstract

The global dissemination of Enterobacterales producing both metallo-β-lactamases (MBLs) and serine β-lactamases (SBLs) represents a critical threat to modern medicine, as no currently marketed antibiotics effectively target MBL-mediated resistance. APC148 is a novel, selective zinc-chelating MBL inhibitor designed to restore β-lactam activity in MBL positive isolates, when used in combination with a broad-spectrum carbapenem. In this study, we evaluated the *in vitro* efficacy of APC148 in triple combinations with either meropenem-avibactam (APC301) or cefepime-avibactam (APC302) against a diverse global collection (JMI collection) of 176 MBL- and SBL-producing Enterobacterales isolates (including NDM, VIM, and IMP variants). Using broth microdilution, the triple combinations were compared against several newly approved and late-stage pipeline antibiotic products. Both APC301 and APC302 demonstrated superior potency, achieving a MIC_90_ of 0.12 µg/mL. When applying CLSI breakpoint interpretive criteria for the parent β-lactams, 99.4% of the MBL and SBL-containing isolates were susceptible to APC301, while 97.2% were susceptible to APC302. These results indicate that the addition of a selective MBL inhibitor to an SBL-inhibitor/β-lactam antibiotic effectively bypasses complex co-existing β-lactam resistance mechanisms in multidrug-resistant (MDR) pathogens. Given that MDR Enterobacterales frequently harbor multiple β-lactamase classes simultaneously, these triple combinations constitute a highly promising clinical strategy to address the therapeutic void in MBL-mediated resistance

## Introduction

The global increase in antimicrobial resistance is currently undermining our ability to treat bacterial infections. A cornerstone in the treatment of serious and life-threatening infections caused by multidrug-resistant (MDR) Gram-negative bacterial pathogens such as *Klebsiella pneumoniae* and *Escherichia coli* has been the carbapenem β-lactam antibiotics. The major advantage of carbapenems has been their relative resistance to β-lactamases (BLs), such as the extended-spectrum β-lactamases (ESBLs) and AmpCs, which constitute common resistance mechanisms against β-lactams [1]. β-lactamases are ancient enzymes produced by bacteria to hydrolyze the β-lactam ring in β-lactams antibiotics. They can be divided into two main types: the serine β-lactamases (SBLs) which utilize an active site serine for hydrolysis and metallo-β-lactamases (MBLs) which require divalent zinc ions present in the active site for the same process [2].

At present, a global increase is observed with respect to the dissemination and diversity of BLs with the ability to inactivate carbapenems (carbapenemases) [3]. The concurrent impact of carbapenem-resistance is illustrated in a European study in which carbapenem-resistance is a significant contributor to the burden of infections caused by antibiotic-resistant bacteria across multiple countries [4]. The introduction of new, more potent derivatives of existing antibiotics provides temporary solutions, however it is not unusual for existing resistance mechanisms to rapidly adapt to accommodate the new derivatives [5], creating a need for continuous development of new treatment options alongside other strategies such as antibiotic stewardship, improved hygiene measures and others.

The introduction of serine β-lactamase inhibitors (SBLi) such as avibactam, vaborbactam and relebactam used in combination with β-lactams has provided treatment options against serine carbapenemase-producing Gram-negative pathogens [6, 7]. However, none of these β-lactamase inhibitors possess inhibitory activity against MBLs.

Consequently, new treatment options for infections caused by MBL-producing Gram-negatives, including NDM-producing *Enterobacterales*, are urgently required. Investigational compounds exist, such as the two cyclic boronate esters taniborbactam and xeruborbactam [8, 9], and the two diaza-bicycooctanes nacubactam and zidebactam, which have shown variable activity in combination with carbapenem antibiotics against MBL-producers [10, 11]. No selective and efficient MBL inhibitors are, however, currently approved for clinical use.

The therapeutic potential of the APC148-carbapenem combination against MBL-producing Gram-negative pathogens has been described earlier [12]. APC148 is a highly promising MBL inhibitor, capable of operating in a functional space not presently filled by any clinically approved compound today. At the present time, APC148 is undergoing clinical development. A first-in-human study in healthy volunteers has been successfully completed. APC148 was well tolerated, and achieved systemic exposures considered suitable for combination therapy [13]. Recently, increased frequencies of patient isolates carrying both MBL and SBL enzymes have been observed. Here, we show that the combined use of APC148, an SBLi and a β-lactam antibiotic in triple combinations demonstrate overall excellent efficiency in lowering meropenem MIC values below clinical breakpoint when tested against a large panel of MDR Enterobacterales simultaneously carrying genes encoding MBL and SBL enzymes.

## Results

The present study assessed the *in vitro* antimicrobial activity of the combinations of APC148 (fixed at 16 µg/mL) with meropenem-avibactam (fixed at 8 µg/mL) (APC301) and cefepime-avibactam (fixed 8 at µg/mL) (APC302), respectively, against 176 MBL-producing Enterobacterales isolates for which detailed information on β-lactamase gene content is available. The two products were compared to commonly used combination products (antibiotic + SBLi) (aztreonam-avibactam (fixed at 4µg/mL) and ceftazidime-avibactam (fixed at 4 µg/mL)) and new pipeline products aztreonam-nacubactam (1:1), cefepime-nacubactam (1:1), cefepime-taniborbactam (fixed at 4 µg/ml) as well as meropenem alone.

### Determination of MIC for APC301 and APC302 Against 176 Enterobacterales Strains

Antimicrobial activities for APC301 and APC302and selected comparator agents are presented as MIC frequency and cumulative percent inhibition in Table 1 for the Enterobacterales isolates combined, includingthe *E. coli* and *K. pneumoniae* isolates. Comprehensive data for all antibiotics and antibiotic/SBLi combinations for all isolates are found in Supplementary Tables S1-S5. Similarly, results presenting MIC range, MIC_50_, MIC_90_, and percent susceptible, intermediate, and resistant according to CLSI (2024) and EUCAST (2024) breakpoint interpretive criteria [14, 15] for selected comparative agents are presented in Table 2 for the Enterobacterales isolates combined including *E. coli* and *K. pneumoniae* isolates, and in Supplementary Tables S6-S10 for all antibiotics for all isolates. Historical MIC data from the SENTRY Antimicrobial Surveillance Program was included in the tables, for: amikacin, aztreonam, cefepime, ceftazidime, ciprofloxacin, colistin, imipenem, and piperacillin-tazobactam (as available). Meropenem was used as the bridge compound between the SENTRY results and the results of this study.

**Table 1.**
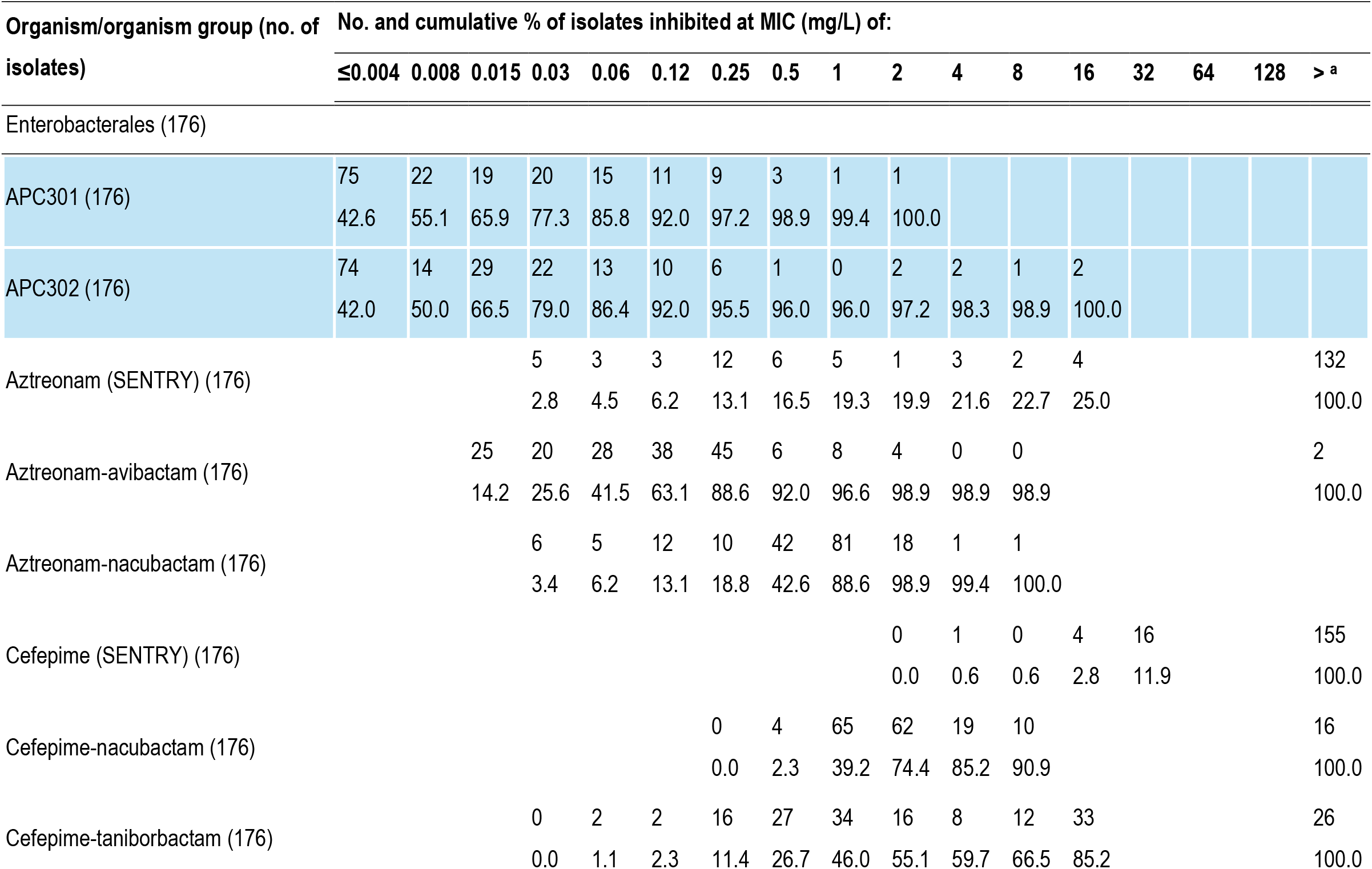
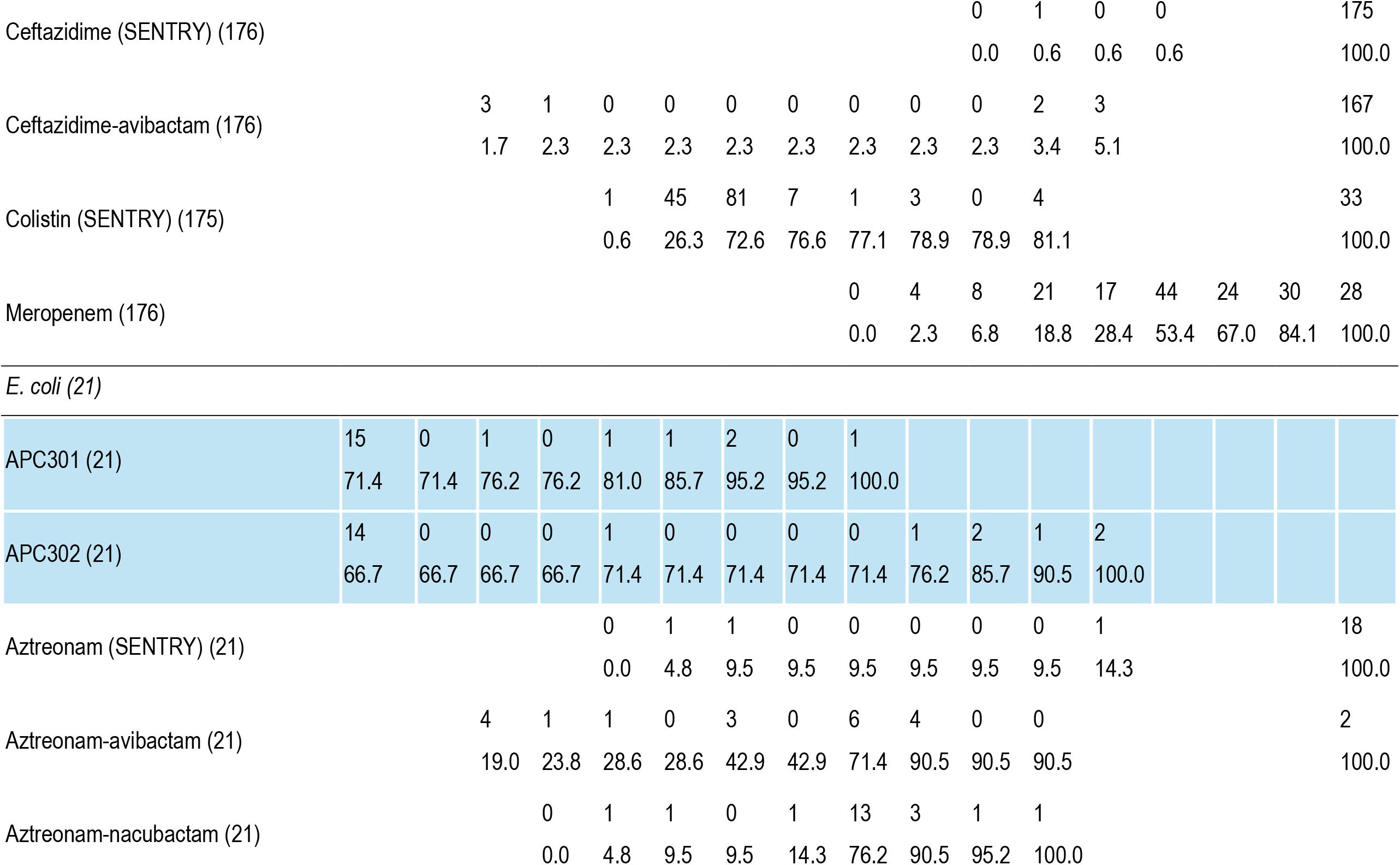
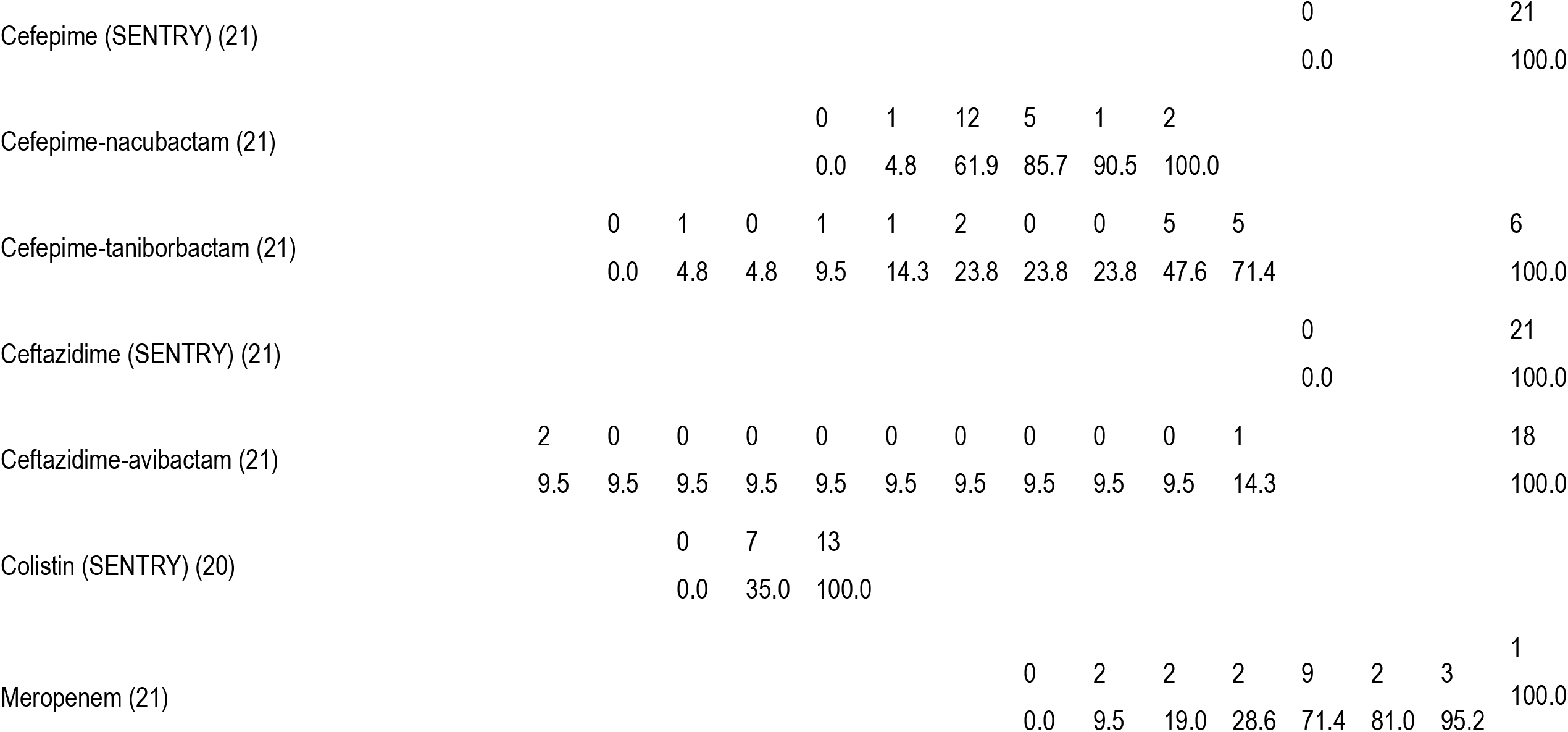
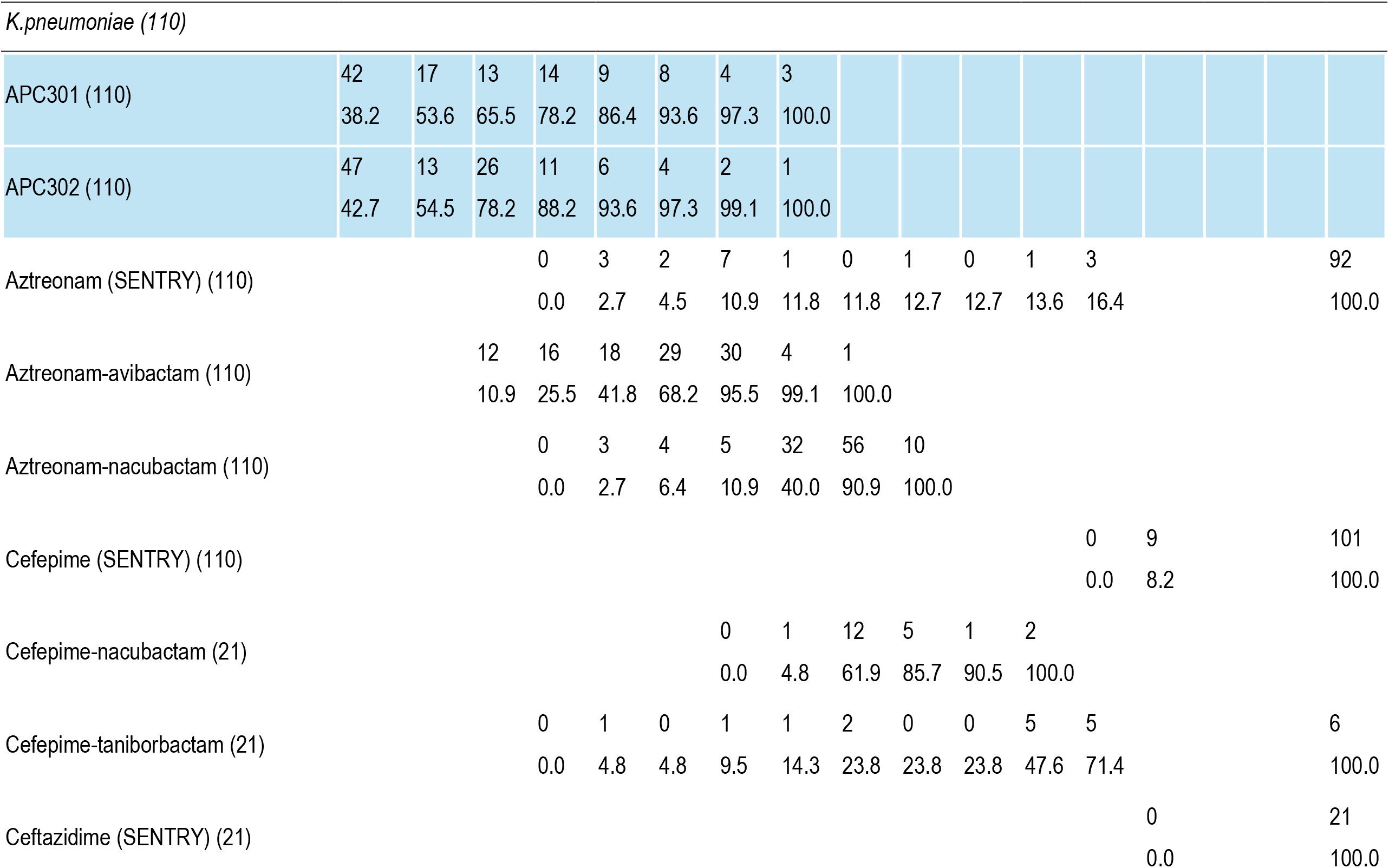
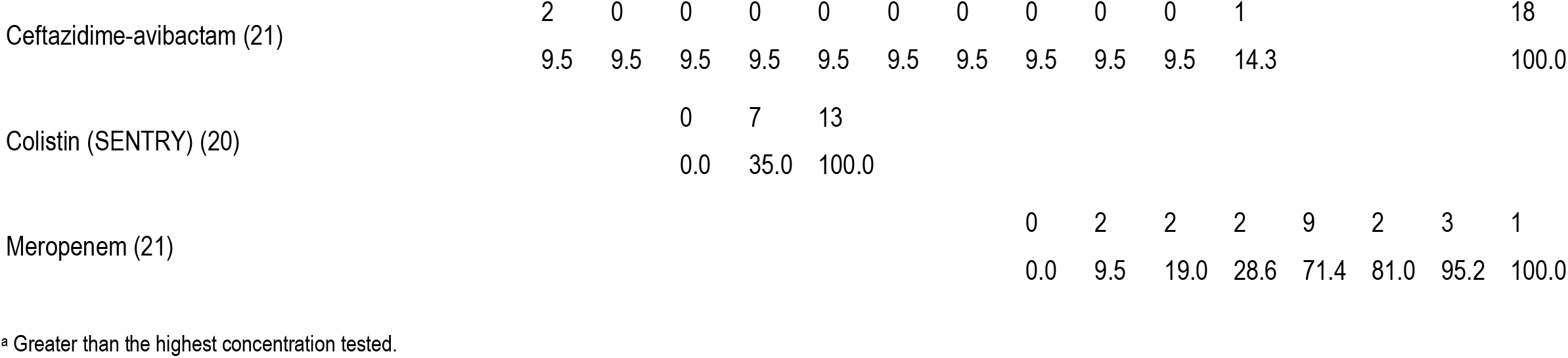
MIC frequency and cumulative percent inhibition for APC301 and APC302 and comparator agents against MBL producing Enterobacterales, including E. coli and K. pneumoniae isolates.

**Table 2.** Antimicrobial activity of APC301 and APC302 and comparator agents tested against Enterobacterales, including E. coli and K. pneumoniae isolates.

| Organism/organism group<br>(no. of isolates) | mg/L |  |  | CLSI<sup>a</sup> |  |  | EUCAST<sup>a</sup> |  |  |
| --- | --- | --- | --- | --- | --- | --- | --- | --- | --- |
|  | MIC<sub>50</sub> | MIC<sub>90</sub> | MIC range | %S | %I | %R | %S | %I | %R |
| *Enterobacterales*<sup>h</sup> (176) |  |  |  |  |  |  |  |  |  |
| APC301 (176) | 0.008 | 0.12 | ≤0.004 to 2 |  |  |  |  |  |  |
| APC302 (176) | 0.008 | 0.12 | ≤0.004 to 16 |  |  |  |  |  |  |
| Aztreonam (SENTRY) (176) | >16 | >16 | ≤0.03 to >16 | 21.6 | 1.1 | 77.3 | 19.3 | 2.3 | 78.4 |
| Aztreonam-avibactam (176) | 0.12 | 0.5 | ≤0.015 to >8 |  |  |  | 98.9 |  | 1.1 |
| Aztreonam-nacubactam (176) | 1 | 2 | ≤0.03 to 8 |  |  |  |  |  |  |
| Cefepime (SENTRY) (176) | >32 | >32 | 4 to >32 | 0.0<sup>d</sup> | 0.6 | 99.4 | 0.0 | 0.6 | 99.4 |
| Cefepime-nacubactam (176) | 2 | 8 | 0.5 to >8 |  |  |  |  |  |  |
| Cefepime-taniborbactam (176) | 2 | >16 | 0.06 to >16 |  |  |  |  |  |  |
| Ceftazidime (SENTRY) (176) | >32 | >32 | 8 to >32 | 0.0 | 0.6 | 99.4 | 0.0 | 0.0 | 100.0 |
| Ceftazidime-avibactam (176) | >16 | >16 | ≤0.015 to >16 | 3.4 |  | 96.6 | 3.4 |  | 96.6 |
| Colistin (SENTRY) (175) | 0.25 | >8 | ≤0.06 to >8 |  | 78.9 | 21.1 | 78.9 |  | 21.1 |
| Meropenem (176) | 32 | >128 | 2 to >128 | 0.0 | 2.3 | 97.7 | 2.3<sup>f</sup> |  | 97.7 |
|  |  |  |  |  |  |  | 2.3<sup>g</sup> | 16.5 | 81.2 |
| *E. coli* (21) |  |  |  |  |  |  |  |  |  |
| APC301 (21) | ≤0.004 | 0.25 | ≤0.004 to 1 |  |  |  |  |  |  |
| APC302 (21) | ≤0.004 | 8 | ≤0.004 to 16 |  |  |  |  |  |  |
| Aztreonam (SENTRY) (21) | >16 | >16 | 0.12 to >16 | 9.5 | 0.0 | 90.5 | 9.5 | 0.0 | 90.5 |
| Aztreonam-avibactam (21) | 1 | 2 | ≤0.015 to >8 |  |  |  | 90.5 |  | 9.5 |
| Aztreonam-nacubactam (21) | 1 | 2 | 0.06 to 8 |  |  |  |  |  |  |
| Cefepime (SENTRY) (21) | >32 | >32 | >32 to >32 | 0.0<sup>d</sup> | 0.0 | 100.0 | 0.0 | 0.0 | 100.0 |
| Cefepime-nacubactam (21) | 1 | 4 | 0.5 to 8 |  |  |  |  |  |  |
| Cefepime-taniborbactam (21) | 16 | >16 | 0.06 to >16 |  |  |  |  |  |  |
| Ceftazidime (SENTRY) (21) | >32 | >32 | >32 to >32 | 0.0 | 0.0 | 100.0 | 0.0 | 0.0 | 100.0 |
| Ceftazidime-avibactam (21) | >16 | >16 | ≤0.015 to >16 | 9.5 |  | 90.5 | 9.5 |  | 90.5 |
|  | MIC<sub>50</sub> | MIC<sub>90</sub> | MIC range | %S | %I | %R | %S | %I | %R |
| Colistin (SENTRY) (21) | 0.25 | 0.25 | 0.12 to 0.25 |  | 100.0 | 0.0 | 100.0 |  | 0.0 |
| Meropenem (21) | 32 | 128 | 4 to >128 | 0.0 | 0.0 | 100.0 | 0.0<sup>f</sup> |  | 100.0 |
|  |  |  |  |  |  |  | 0.0<sup>g</sup> | 19.0 | 81.0 |
| *K. pneumoniae* (110) |  |  |  |  |  |  |  |  |  |
| APC301 (110) | 0.008 | 0.12 | ≤0.004 to 0.5 |  |  |  |  |  |  |
| APC302 (110) | 0.008 | 0.06 | ≤0.004 to 0.5 |  |  |  |  |  |  |
| Aztreonam (SENTRY) (110) | >16 | >16 | 0.06 to >16 | 12.7 | 0.9 | 86.4 | 11.8 | 0.9 | 87.3 |
| Aztreonam-avibactam (110) | 0.12 | 0.25 | ≤0.015 to 1 |  |  |  | 100.0 |  | 0.0 |
| Aztreonam-nacubactam (110) | 1 | 1 | 0.06 to 2 |  |  |  |  |  |  |
| Cefepime (SENTRY) (110) | >32 | >32 | 32 to >32 | 0.0<sup>d</sup> | 0.0 | 100.0 | 0.0 | 0.0 | 100.0 |
| Cefepime-nacubactam (110) | 2 | 8 | 0.5 to >8 |  |  |  |  |  |  |
| Cefepime-taniborbactam (110) | 2 | >16 | 0.12 to >16 |  |  |  |  |  |  |
| Ceftazidime (SENTRY) (110) | >32 | >32 | >32 to >32 | 0.0 | 0.0 | 100.0 | 0.0 | 0.0 | 100.0 |
| Ceftazidime-avibactam (110) | >16 | >16 | ≤0.015 to >16 | 2.7 |  | 97.3 | 2.7 |  | 97.3 |
| Colistin (SENTRY) (110) | 0.25 | >8 | 0.12 to >8 |  | 79.1 | 20.9 | 79.1 |  | 20.9 |
| Meropenem (110) | 64 | >128 | 2 to >128 | 0.0 | 0.9 | 99.1 | 0.9<sup>f</sup> |  | 99.1 |
|  |  |  |  |  |  |  | 0.9<sup>g</sup> | 11.8 | 87.3 |

<sup>a</sup> Criteria as published by CLSI (2024) and EUCAST (2024). S: Susceptible, I: Intermediate / Susceptible, increased exposure, R: Resistant
<sup>b</sup> For infections originating from the urinary tract. For systemic infections, aminoglycosides must be used in combination with other active therapy.
<sup>c</sup> CLSI M100 standard is recognized.
<sup>d</sup> Intermediate is interpreted as susceptible-dose dependent.
<sup>e</sup> CLSI M100 standard is recognized. Intermediate is interpreted as susceptible-dose dependent.
<sup>f</sup> Using meningitis breakpoints.
<sup>g</sup> Using non-meningitis breakpoints
<sup>h</sup>Organisms include *Citrobacter freundii* species complex (3), *Enterobacter cloacae* species complex (17), *Escherichia coli* (21), *Klebsiella aerogenes* (1), *K. oxytoca* (14), *K. pneumoniae* (110), *Proteus mirabilis* (4), *Providencia rettgeri* (3), *P. stuartii* (2), and *Serratia marcescens* (1).

### Antimicrobial Activity of APC301 and APC320 and Comparator Agents against

#### Enterobacterales Isolates

Of all the tested BL and BLI combinations in this study, APC301 and APC302 were the most potent combinations tested against a collection of 176 MBL containing Enterobacterales isolates, with MIC_50/90_ values of 0.008/0.12 µg/mL. The other most effective combinations included aztreonam-avibactam (MIC_50/90_, 0.12/0.5 µg/mL) and aztreonam-nacubactam (MIC_50/90_, 1/2 µg/mL) (Tables 2 and S6). Susceptibility of the MBL containing Enterobacterales isolates ranged from 0.0% (cefepime, ceftazidime, imipenem, and meropenem) to 39.8% (amikacin) using CLSI breakpoint criteria; 0.0% (cefepime, ceftazidime, and imipenem) to 98.9% (aztreonam-avibactam) using EUCAST breakpoint criteria (Tables 2 and S6). Applying CLSI breakpoint interpretive criteria for cefepime and meropenem to the APC combinations, 99,4% of the MBL containing Enterobacterales isolates would be susceptible to APC301 and 97.2% would be susceptible to APC302 (Table 1 and S1).

#### Escherichia coli Isolates

When measuring the antimicrobial activity of the BL and BLI combinations, APC301 was the most active against a collection of 21 MBL containing *E. coli* isolates, with MIC_50/90_ values of ≤0.004/0.25 µg/mL. Aztreonam-avibactam and aztreonam-nacubactam were also effective, with MIC50/90 values of 1/2 µg/mL) (Tables 2 and S7).

Susceptibility of the MBL containing *E. coli* isolates ranged from 0.0% (cefepime, ceftazidime, imipenem, meropenem, and piperacillin-tazobactam) to 52.4% (amikacin) using CLSI breakpoint criteria; 0.0% (cefepime, ceftazidime, imipenem, and piperacillin-tazobactam) to 100.0% (colistin) using EUCAST breakpoint criteria (Tables 2 and S7).

Applying CLSI breakpoint interpretive criteria for cefepime and meropenem to the APC combinations, 76.2% of the MBL containing *E. coli* isolates would be susceptible to APC302 and 100.0% would be susceptible to APC301 (Tables 1 and S2).

#### Klebsiella pneumoniae Isolates

APC301 and APC302 were the most active combinations tested against a collection of 110 MBL containing *K. pneumoniae* isolates with MIC50/90 values of 0.008/0.06 µg/mL and 0.008/0.12 µg/mL, respectively. Aztreonam-avibactam and aztreonam-nacubactam were also active with MIC_50/90_ values of 0.12/0.25 µg/mL and 1/1 µg/mL, respectively (Tables 2 and S8).

Susceptibility of the MBL containing *K. pneumoniae* isolates ranged from 0.0% (cefepime, ceftazidime, imipenem, meropenem, and piperacillin-tazobactam) to 33.6% (amikacin) using CLSI breakpoint criteria; 0.0% (cefepime, ceftazidime, imipenem, and piperacillin-tazobactam) to 100.0% (aztreonam-avibactam) using EUCAST breakpoint criteria (Table 2 and S8).

Applying CLSI breakpoint interpretive criteria for cefepime and meropenem (alone) to the APC combinations, 100.0% of the MBL containing *K. pneumoniae* isolates would be susceptible to APC301 and to APC302 (Table 1 and S3).

#### Enterobacter cloacae species complex Isolates

APC301, APC302 and aztreonam-avibactam were the most active combinations tested against a collection of 17 MBL containing *E. cloacae* species complex isolates, with MIC_50/90_ values of 0.06/0.25 µg/mL, 0.008/0.03 µg/mL, and 0.12/0.25 µg/mL, respectively. Aztreonam-nacubactam was also effective, with an MIC_50/90_ value of 0.5/1 µg/mL) (Table S9).

Susceptibility of the MBL containing *E. cloacae* species complex isolates ranged from 0.0% (cefepime, ceftazidime, ceftazidime-avibactam, imipenem, meropenem, and piperacillin-tazobactam) to 52.9% (amikacin and aztreonam, respectively) using CLSI breakpoint interpretive criteria; 0.0% (cefepime, ceftazidime, ceftazidime-avibactam, imipenem, and piperacillin-tazobactam) to 100.0% (aztreonam-avibactam) using EUCAST breakpoint criteria (Table S9).

Applying CLSI breakpoint interpretive criteria for cefepime and meropenem to the APC combinations, 100.0% of the MBL containing Enterobacterales isolates would be susceptible to APC301 and APC302 (Table S5).

#### Klebsiella oxytoca Isolates

APC301 and APC302 were the most active combinations tested against a collection of 14 MBL containing *K. oxytoca* isolates with MIC_50/90_ values of ≤0.004/0.06 µg/mL and ≤0.004/0.03 µg/mL, respectively. Aztreonam-avibactam was also active with an MIC_50/90_ of 0.12/0.25 µg/mL) (Table S10). Susceptibility of the MBL containing *K. oxytoca* isolates ranged from 0.0% (cefepime, ceftazidime, ceftazidime-avibactam, imipenem, meropenem, and piperacillin-tazobactam) to 28.6% (amikacin) using CLSI breakpoint criteria; 0.0% (cefepime, ceftazidime, ceftazidime-avibactam, imipenem, meropenem, and piperacillin-tazobactam) to 100.0% (aztreonam-avibactam) using EUCAST breakpoint criteria (Table S10).

Applying CLSI breakpoint interpretive criteria for cefepime and meropenem to the APC combinations, 100.0% of the MBL containing *K. oxytoca* isolates would be susceptible both to APC301 and to APC302 (Table S5).

## Discussion

In this study we report the *in vitro* activity of APC301 and APC302 in a collection of 176 MBL producing Enterobacterales isolates. The two products were compared to commonly used, newly approved and late-stage pipeline antibiotics. Of all combinations tested, APC301 and APC302 showed by far the strongest MIC reduction. Nearly all isolates were resistant to meropenem and cefepime alone, and to ceftazidime-avibactam, commonly used antibiotics. Interestingly, also the last-resort antibiotic colistin had a MIC_90_ of >8, rendering most of these isolates resistant according to CLSI and EUCAST breakpoints. While in the present study meropenem and cefepime were combined with APC148 and avibactam at 8 µg/mL, results from a recent study in 303 multidrug-resistant *E. coli* and *K. pneumoniae* (298 MBL-producing) from India, showed that APC301 using avibactam at 4 µg/mL was as equally as effective as avibactam 8 µg/mL [16].

Cefepime combined with new SBL-inhibitors like taniborbactam and nacubactam performed poorly with MIC_90_s of 16 and 8 µg/mL respectively. These SBL-inhibitor products have shown promise towards some MBL-producing isolates but were clearly inferior in this collection, most likely due to variable inhibition of MBLs.

The MBL-producing isolates were more susceptible towards the aztreonam products. Considering that aztreonam is the only β-lactam antibiotic that is not hydrolyzed by MBLs [17], the potency of these combinations is perhaps not surprising [18]. However, resistance to cephalosporins and monobactams caused by mutations in the PBP receptor is increasingly being reported [19-22]

Overall, 99.4% of the 176 MBL containing Enterobacterales isolates would be considered susceptible to APC301 when CLSI breakpoint interpretive criteria were applied. The single non-susceptible isolate was the one *Serratia marcescens* isolate tested, carrying a VIM-1 MBL. All but 5 of the 176 MBL containing Enterobacterales isolates (97.2%) would be considered susceptible to APC302 when CLSI breakpoint interpretive criteria were applied. The 5 non-susceptible isolates were all *E. coli* isolates containing either an NDM-5 (n=4) or NDM-19 (n=1) MBL (Table S11). Resistance to cephalosporins in strains carrying NDM-5 has previously been shown to be linked to PBP3 mutations [22].

The six Enterobacterales strains carrying IMP variants were all susceptible to APC148 in combination with cefepime/avibactam as well as with meropenem/avibactam. Considering that IMP family MBLs are known to be less susceptible to inhibition by taniborbactam in particular [23], it is interesting to note that for the six Enterobacterales strains harboring an IMP variant, APC301 and APC302 show MIC values ranging between ≤0.04 and 0.25 µg/mL, rendering these the most potent combinations against MBLs (Table S12). In contrast, MIC values for cefepime-taniborbactam falls between 8 and >16 µg/mL (Table S11).

In summary, this study highlights the high efficacy of the *in vitro* activity of APC301 and APC302 against a wide range of clinical isolates of MBL containing Enterobacterales and supports the continued evaluation of these antibacterial combinations.

Considering that multi-resistant Enterobacterales commonly produce multiple β-lactamases, including combinations of both SBLs and MBLs, a selective MBL inhibitor such as APC148 when combined with an SBLi and a carbapenem antibiotic (APC301 and APC302), could constitute a novel and clinically promising treatment option, in particular in light of emerging resistance observed with combinations with aztreonam. The high efficacy observed in the present study, together with *in vivo* efficacy data and the good tolerability of APC148 demonstrated in a phase 1 study, suggests a promising combination therapy to address the unmet medical need for treatment against multidrug resistant bacteria.

## Materials and Methods

### Bacterial strains used in this study

A complete list of the 176 Enterobacterales strains tested are listed in Table S13 along with molecular characterization of β-lactamases from prior analysis. Each of the Enterobacterales isolates tested contained at least one MBL gene. In addition to the MBL, most isolates also carried one or more SBL genes. The bacterial strains tested were from the JMI collection (www.jmilabs.com) and included Clinical and Laboratory Standards Institute reference and quality control (QC) strains as well as recent clinical isolates from the following organism groups: *Enterobacter cloacae* species complex, *Escherichia coli, Klebsiella oxytoca*, and *Klebsiella pneumoniae*. Bacterial identifications were confirmed by Element Iowa City (JMI Laboratories) using matrix-assisted laser desorption ionization-time of flight mass spectrometry (MALDI-TF MS) (Bruker Daltonics, Bremen, Germany).

### Bacterial Isolates Geographic Regions

Isolates were collected in 49 medical centers located in 24 countries and 8 U.S. Census divisions: the U.S. (16 medical centers; 23 isolates; 13.1% of overall isolates), Europe (21 medical centers; 89 isolates; 50.6% overall), Latin America (5 medical centers; 18 isolates; 10.2% overall), and the Asia-Pacific region (7 medical centers; 46 isolates; 26.1% overall) between 2019-2022, as part of the SENTRY Antimicrobial Surveillance Program.

### Bacterial Isolate Infection Types

The isolates were collected primarily from patients with bloodstream infections (47 isolates; 26.7% overall), skin and skin structure infections (26 isolates; 14.8% overall), pneumonia in hospitalized patients (52 isolates; 29.5% overall), urinary tract infections (37 isolates; 21.0% overall), intraabdominal infections (8 isolates; 4.5% overall), and other infection types (6 isolates; 3.4% overall), according to a common surveillance design.

## Susceptibility Testing Methods

### Broth Microdilution

Enterobacterales isolates were tested for antimicrobial susceptibility using broth microdilution susceptibility panels according to CLSI M07 (2025) and M100 (2025) guidelines [14, 24], using Element Iowa City produced frozen-form, 96-well MIC panels. The testing medium was cation-adjusted Mueller-Hinton broth (Becton Dickinson, Franklin Lakes, NJ USA). The zinc concentration of this medium was measured to 0.92 µg/mL (Eurofins (Test America) IL, USA). Single MIC values were measured for the compounds and combinations tested against each of the 176 bacterial isolates and QC strains tested. The MICs of meropenem and cefepime were tested with avibactam at a fixed value of 8 µg/mL and APC148 at 16 µg/mL throughout the study, unless otherwise stated.

### Antimicrobial Compounds and Enzyme inhibitors

APC148 and comparator agent compound information, including potency, range tested, solvent, and diluent are presented in Table S12.

### Interpretive Criteria

All categorical interpretations for comparator agents used CLSI M100 (2024) or EUCAST v14.0 (2024) breakpoint criteria [14, 15, 25]

## Quality Control

Element Iowa City (JMI Laboratories) followed current CLSI quality assurance practices when performing the susceptibility tests. MIC values were validated by concurrently testing QC and reference strains from ATCC, as recommended by CLSI (M100, 2024) [14]. MIC values were validated by concurrently testing control agents (aztreonam-avibactam, aztreonam-nacubactam, cefepime-nacubactam, cefepime-taniborbactam, ceftazidime-avibactam and meropenem alone) against CLSI-recommended QC strains including: *E*.*coli* NCTC 13353, *K. pneumoniae* ATCC 700603, *K. pneumoniae* ATCC BAA-1705, *K. pneumoniae* ATCC BAA-2814, and *P. aeruginosa* ATCC 27853. The initial inoculum density (target: 5 x 10^5^ CFU/mL) during susceptibility testing was monitored by bacterial colony counts (live cell counts, CFU counts).

No out-of-range QC results were observed in this study.

## ACKNOWLEDGEMENTS

This work was funded through the IPN programme at The Research Council of Norway (project number 346300), and the support is gratefully acknowledged. The funders had no role in study design, data collection or interpretation, nor in the decision to submit the work for publication.

